# Design and Characterisation of a Sensor Integrating Chemical and Temperature Inputs for Gene Expression

**DOI:** 10.1101/2025.07.24.666488

**Authors:** V. Kumar, S. Sen

## Abstract

The modularity of transcription factors has facilitated the development of combinatorial promoters that regulate gene expression in response to chemical inducers. Nevertheless, controlling gene expression by the joint influence of a chemical inducer and an environmental variable, such as temperature, has not been as extensively studied or analyzed. We designed a two-input combinatorial sensor combining the action of a chemical inducer (Anhydrotetracycline, aTc) and temperature. The design was based on transcriptional regulation by the *P*_*tet*_ promoter and translational regulation by an RNA thermometer. The design functioned as an AND Gate, exhibiting a four-fold change between a completely ON state, with aTc at 37 °C, and a completely OFF state, without aTc at 29 °C. We mapped the design into a Logic-Symmetry space and found that the design functioned like an AND Gate with an asymmetry of 0.266. This work should contribute to developing a temperature-based combinatorial regulation system.

## 1. Introduction

A dominant workflow in synthetic biology is to use building blocks from nature to generate new regulatory devices with novel functionality. RNA-based genetic circuits have emerged as an attractive substrate for biomolecular regulation due to their modularity and versatility [1].

A key aspect of biomolecular regulation is the presence of combinatorial controls, which are naturally prevalent in biological systems. An example of this is in the regulation of a Lac operon in the presence of CAP and lactose [2], [3]. Building on this concept, studies have demonstrated how the modular nature of RNA thermometers can be harnessed for combinatorial regulation in conjunction with riboswitches [4], [5]. Here, the ligand binding and the temperature sensing are utilized as triggers. In a recent study, a chemical inducer (IPTG) and temperature were used in a combinatorial fashion to respond to heat in protocells [6]. A prerequisite for designing combinatorial sensors from natural regulators is that the combined building blocks should be sufficiently modular and functional in different genetic contexts. This relies on the modularity of regulation in multiple layers to design different phenotypes in the cell. Thus, the modular nature of RNA thermometers can be potentially exploited with a different chemical inducer to regulate gene expression.

In this work, we aimed to design a sensor that integrates chemical inducer (aTc) and temperature inputs in a combinatorial fashion. We used Green fluorescence protein (GFP) as a reporter and estimated the sensor output using measurements of cell density and fluorescence in growing cells. The design combined the chemical and temperature inputs using a *P*_*tet*_ promoter driving an RNA thermometer so that transcription and translation could be regulated by aTc and temperature, respectively. We found that the combinatorial sensor exhibited a four-fold change between a completely ON state, with aTc and at 37 °C, and a completely OFF state, without aTc and at 29 °C. We mapped the measurements into a logic-symmetry space and found that the design functioned like an AND Gate with an asymmetry of 0.266.

Further, in designs without the *P*_*tet*_ promoter and/or the RNA thermometer, the output remained relatively the same, depending on the aTc level and temperature. The experimental results aligned with expectations, demonstrating the effectiveness of the design strategy. This dual-input sensor system may be useful for synthetic biology applications.

## II. Results and Discussion

### A Sensor Design

Combinatorial sensor circuits, such as for an AND Gate, involve the detection and processing of multiple input signals. A general design strategy is to utilize modular genetic components, such as promoters, transcription factors, and ribosome binding sites [7].

The basic working principle of the sensor is the integration of a chemical (aTc) and temperature inputs through a *P*_*tet*_/TetR system with an RNA thermometer regulating the Ribosome Binding Site. We designed the sensor using a combinatorial promoter driving the expression of GFP. The promoter of GFP was chosen to be *P*_*tet*_, which can be regulated by the aTc inducer. The thermometer *U* 2 [8] is inserted downstream of the promoter *P*_*tet*_. In this circuit, the presence of aTc prevents the suppression of TetR. This design is expected to express GFP more in the presence of both aTc and temperature than when either is absent. The design acts on the transcription and translation in a sequential manner and is anticipated to function as an AND logic circuit.

The experimental construct for the design was encoded on a plasmid (Supplementary Figure 1). The plasmid contains the promoter, the RNA thermometer, and GFP as a reporter gene, flanked by terminators. We constructed controls for the action of temperature and for the action of the aTc inducer. A constitutive promoter *P*_*rrn*_ was used instead of *P*_*tet*_ to control for the effect of aTc (Fig. 1(b)). This control is expected to not undergo a substantial change depending on the aTc level. To control for the effect of temperature, an RNA thermometer *U* 0 [8] that is believed to not respond to temperature was used instead of *U* 2 (Fig. 1(c)). A fourth construct that combines both the above controls was also used (Fig. 1(d)).

**Fig. 1.**
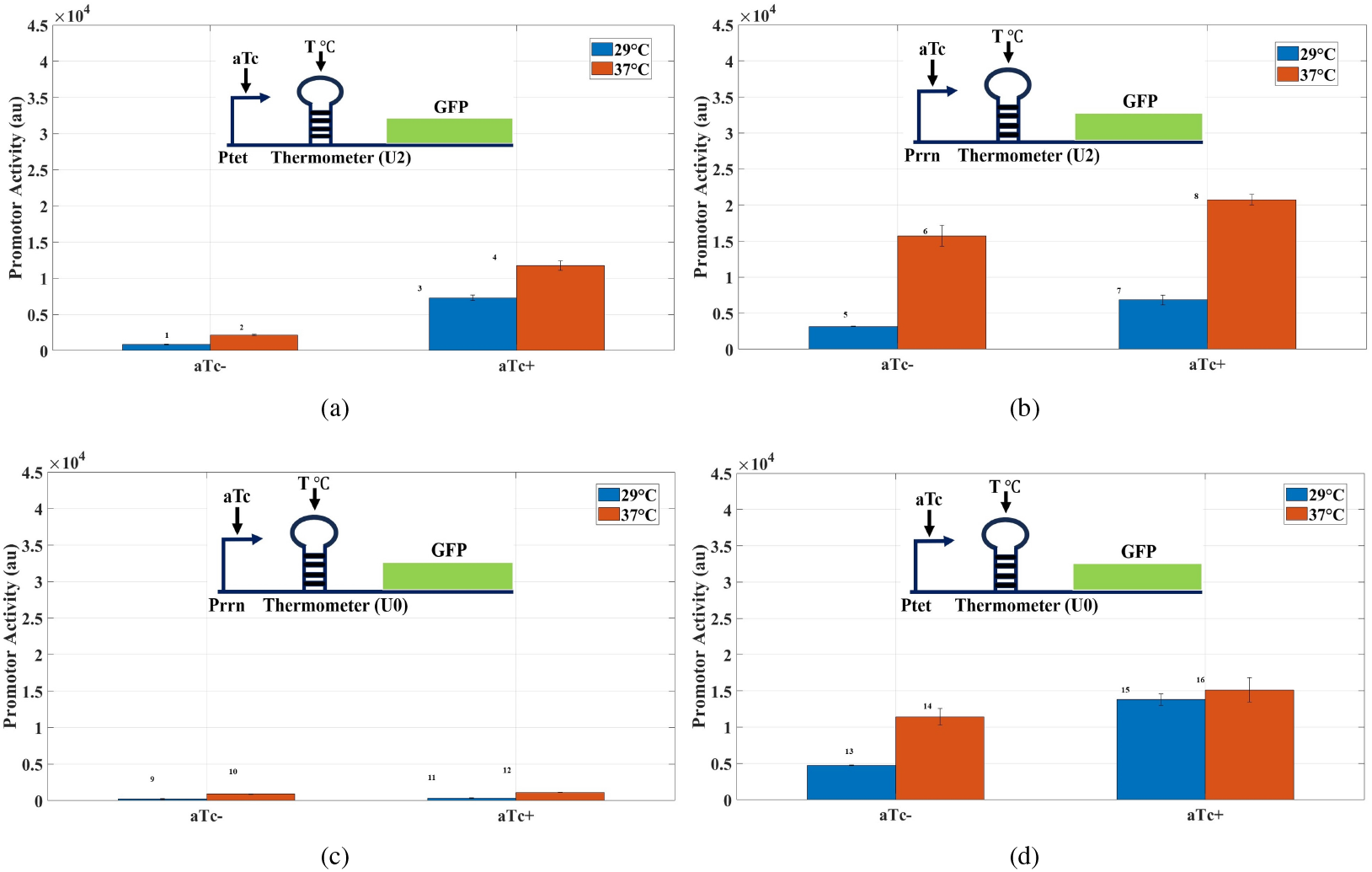
Combinatorial sensor. Blue and red bars represent the promoter activities at 29 °C and 37 °C, respectively, for constructs with (a) Promoter *P*_*tet*_ and RNA thermometer (*U* 2), (b) Constitutive promoter *P*_*rrn*_ and RNA thermometer (*U* 2), (c) Consecutive and *U* 0 (No thermometer or Weak thermometer, and (d) Promoter *P*_*tet*_ and weak RNA thermometer (*U* 0). Promoter activity (*α*) was calculated at the 250^th^ minute after aTc induction for all circuits representing the midpoint of the measurement window. Promoter activity at 100^th^, and 350^th^ minutes are in Supplementary Figures 5 and 6, respectively. The bar heights are a mean of 3 replicates, and the standard deviations are represented as the error bars.

### B. Sensor Characterisation

The dual-input RNA sensor for gene regulation was characterized using a platereader. The detailed experimental protocols and mathematical analysis are described in the Methods section.

The experimental measurements were in line with our expectations. The outcomes of the sensor design are illustrated in (Fig. 1(a)). In this circuit, there was low expression in the absence of aTc, because TetR repressed the promoter. This is consistent with our expectations, as the promoter did not turn on. The presence of aTc prevents the repression by TetR and allows expression from the *P*_*tet*_ promoter. An increase in temperature allows for even higher expression. Thus, in the presence of aTc and at the high temperature (37 °C), the promoter activity is higher than when either is absent.

Constructs with a constitutive promoter and/ or a weak thermometer were used as controls (Figs. 1(b)-(d)). With a constitutive promoter driving an RNA thermometer, the response was primarily like that of an RNA thermometer (Fig. 1(b)). A constitutive promoter driving a weak thermometer did showed low fluorescence levels (Fig. 1 (c)). The weak RNA thermometer driven by the *P*_*tet*_ promoter showed an OR Gate-like behaviour where either input was sufficient to drive high fluorescence. The raw data for optical density (OD), fluorescence (F), and F/OD are in Supplementary Figure 3.

The comparison also demonstrated that the temperature effect in the designed AND logic is primarily dominated by the aTc input. This can be seen in (Fig. 1(a)) as the change in activity due to change in aTc induction at 37 °C is much higher with respect to the change in the activity due to temperature change. This is also evidenced by the GFP (Green fluorescence protein)/ OD (optical density) plot in Supplementary Figure 4 (b). The GFP/OD plots for the other three circuits are also shown in Supplementary Figure 4 (a),(c),(d).

We noted that temperature could affect the secondary structure of the RNA thermometer as well as other aspects of the measurement process, such as the fluorescence parameters of GFP, binding properties of aTc, and possibly the underlying dynamics. However, these factors should affect all the designs in a similar manner. We focused on the relative difference in the sensor activities.

We used a logic-symmetry space to characterize the behaviour of such combinatorial circuits [9]. The promoter activity was mapped to a 2-D space defined by two parameters, logic type (*l*: 0 = OR-like, 1 = AND-like) and asymmetry (*a*: 0 = symmetric response to both inputs, 1 = suggests a single input Gate (SIG-like), where only one input drives the response). Promoter activity (*α*) was measured across four input combinations ([0, 0], [1, 0], [0, 1], [1, 1] for aTc and temperature, respectively) and normalized by basal activity to obtain *α*_*norm*_ (Fig. 2(a)). *α >* 0.7 (High activity), *α <* 0.465 (Low activity) were chosen as thresholds. Logic scores were assigned per input: 0 for [0, 0], 1 for [1, 1], and for [1, 0]/[0, 1], scored as 1 if *α >* 0.7, 0 if *α <* 0.465, or interpolated in between. Constructs were classified as AND-like (high [1, 1], low others), OR-like (high [1, 0]/[0, 1]/[1, 1], low [0, 0]), or complex/additive based on mean of [1, 0]/[0, 1] (Fig. 2(b)). Since the output responses exhibit graded, continuous expression levels across input states, conventional Boolean logic might not capture the full dynamic range and intermediate behaviors inherent to these circuits. If a construct’s *α*_*norm*_ values didn’t fit a predefined logical Gate pattern, it was labeled as complex/additive. Construct points were plotted as (*l, a*) in logic-symmetry space. Asymmetry (*a*) was calculated as 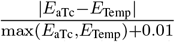, where *E*_aTc_ and *E*_Temp_ are sum of *α*_*norm*_ for high aTc and temperature inputs, respectively. The detailed formulation of parameters *a* and *l* is discussed in the Parameter Formulation Section of the Supplementary Material.

**Fig. 2.**
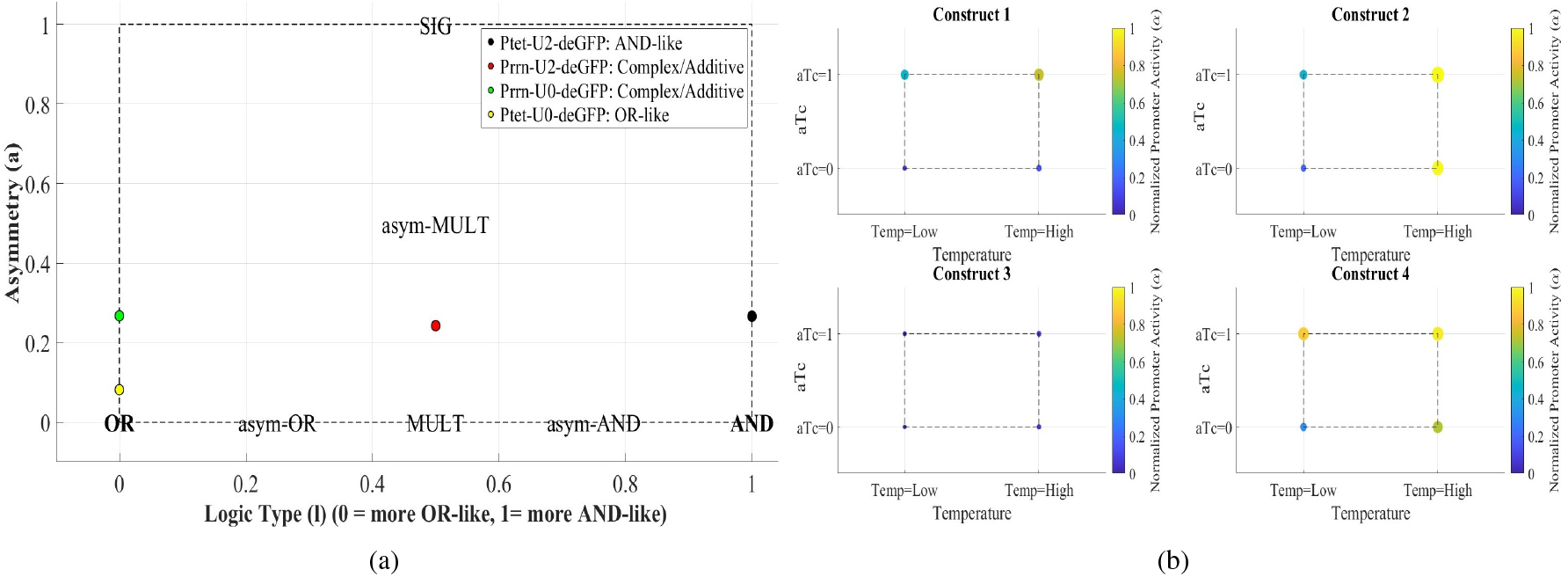
Logic Symmetric space. Logical phenotypes represented are Construct 1: *P*_*tet*_ − *U*2 − *deGFP*, Construct 2: *P*_*rrn*_ − *U* 2 −*deGFP*, Construct 3: *P*_*rrn*_ −*U*0 −*deGFP*, and Construct 4: *P*_*tet*_ −*U*0 −*deGFP*. (a) The domain and locations of identified logic Gates. Promoter activity is mapped by their asymmetry (y-axis), logic type (x-axis). The asym-OR/asym-AND/Mult/ asym-mult and SIG represent an intermediate logic between AND and OR. “MULT” signifies a multiplicative interaction demonstrating symmetry and intermediate logic rather than a linear trend. (b) Activity profile of each construct. The normalized promoter activity (*α*) was displayed by color markers across four input conditions ([0, 0], [0, 1], [1, 0], [1, 1]) of aTc and temperature, respectively, for each of the constructs. Marker size reflects strength, with larger points indicating stronger promoter response. Construct 1 displays AND-like behavior, exhibiting high activity only when both inputs are high, whereas Constructs 2 – 4 reveal diverse responses, highlighting distinct patterns.

The main sensor design *P*_*tet*_ −*U*2 −*deGFP* exhibited AND-like behavior, requiring both inputs for high output. The control *P*_*tet*_ −*U*0 −*deGFP* was OR-like, responding to either input. The other controls *P*_*rrn*_ −*U*2 −*deGFP* and *P*_*rrn*_ −*U*0 −*deGFP* showed complex/additive behaviors with mixed responses. This analysis quantitatively confirmed the main designs’ distinct logical functions and the controls’ intermediate or basal activities. The quantization used relative promoter activities rather than absolute levels, which are less susceptible to changes in growth medium, reporter selection, and other experimental parameters. Thus, this representation offered qualitative and quantitative classification of different logic types.

### C. Discussion

The ability to combine different inputs in a predictable fashion is important in a toolbox of biomolecular components. We demonstrated the design of a combinatorial sensor that combines a temperature input and a chemical input. The temperature input was translationally sensed using an RNA thermometer, the chemical input was transcriptionally sensed using the aTc-TetR-*P*_*tet*_ system, and the combined effect was measured using GFP expression.

The transcription and translation-based AND Gate design we employed also produces different sensor outputs according to different input combinations. We observed that the *P*_*tet*_-based design with aTc at 37 °C also produces a similar four-fold change to when the same design is subjected to 29 °C without aTc. This quantified gene activity was consistent with an AND Gate operation. We also mapped the promoter activity using logic type and symmetry parameters.

The use of combinatorial sensors is a promising approach to expand the range of input signals and tackle real-world challenges through logical computations. Our findings highlight the potential of combinatorial designs using temperature as an input, which can be used for developing programmable cells and complex synthetic systems. The results should help design circuits that utilize temperature as an input.

## III. Materials and Methods

The following section describes the plasmids, the bacterial strains, and the measurement methods used in this study.

### A. Plasmids and Bacterial strains

Four plasmids, all based on the vector backbone of pBSU2 [8], were made using a combination of gene synthesis and restriction enzyme-based cloning. Each insert was flanked by the two terminators TBS7 and Trps16 and expressed the green fluorescent protein deGFP. The promoter was either the constitutive *P*_*rrn*_ promoter or the aTc-inducible *P*_*tet*_ promoter. The thermometer *U* 2 or *U* 0 were used for each of the promoters resulting in four combinations (Table I). The plasmids replicate from the ColE1 origin and are resistant to ampicillin. These four plasmids, as shown in Table I, were transformed into the *E*.*coli* strain MG1655 Z1. The transformed cells were spotted on LB-agar plates containing 100 *µ*g/ml Ampicillin.

**TABLE 1.**
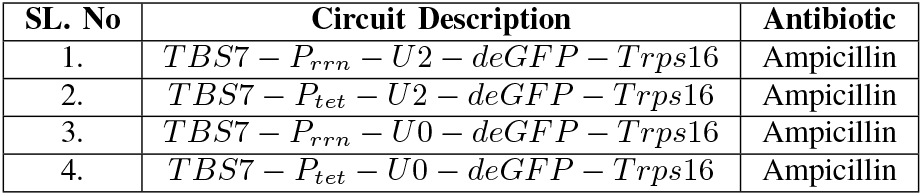
Plasmids description.

| SL. No | Circuit Description | Antibiotic |
| --- | --- | --- |
| 1. | $TBS7 - P_{rrn} - U2 - deGFP - Trps16$ | Ampicillin |
| 2. | $TBS7 - P_{tet} - U2 - deGFP - Trps16$ | Ampicillin |
| 3. | $TBS7 - P_{rrn} - U0 - deGFP - Trps16$ | Ampicillin |
| 4. | $TBS7 - P_{tet} - U0 - deGFP - Trps16$ | Ampicillin |

### B. Measurements

The experiments were done following the process of incubation, streaking inoculation, shaker incubation, and dilution (Supplementary Figure 2). For this, all the bacterial strains were cultured overnight in an LB medium using Ampicillin (100 *µ*g/ml). They were subsequently diluted in the LB media with the ampicillin antibiotic and incubated for two hours at 37 °C and 29 °C. Subsequently, two cultured samples were prepared with aTc concentrations (0 ng/ml and 5 ng/ml) for each strain, and 200 *µ*l of the mixture was transferred onto a 96-well plate (Eppendorf). This well plate containing the bacterial strains was inserted into a microplate reader (Biotek synergy H1) and left to incubate for a duration of 8 hours at temperatures of 37 °C and 29 °C for taking the readings separately. The optical density (absorbance at 600 nm) and fluorescence (excitation/emission = 485/525 nm) of each culture were monitored at 5-minute intervals, with a 2-minute shaking period between each measurement. The measurements were done in triplicate, and the procedure above was repeated for three days at each temperature.

### C. Data Analysis

All the data was analyzed using MATLAB version R2021a. We measured the fluorescence (reflecting the total expression level of the reporter gene GFP in the culture) and the optical density (OD) over time using a plate reader to quantify promoter activity. The samples’ fluorescence and optical density across three days were used to compute the sample mean and standard deviation. The basal absorbance and fluorescence of the background media were subtracted from the sample readings.

We used the following mathematical methodology to calculate the amount of promoter activity. Let *N* represent the total number of bacterial cells and *x* denote the amount of GFP per cell. The *Nx* represents the fluorescence F that changes over time, and *N* can be determined from the optical density (OD) readings recorded.

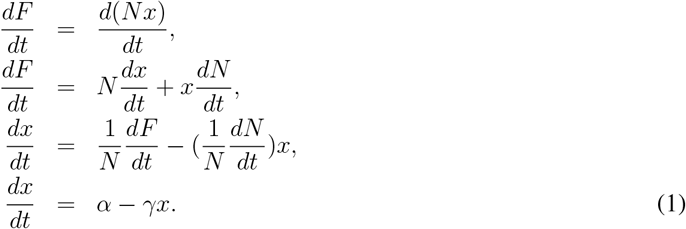

Here, the metric *α* is the production factor, which indicates the rate at which the gene’s promoter starts transcription and is used to represent promoter activity. The *γ* represents the decay or dissolution rate. In the analysis, *γ* was assumed to be dominated by dilution due to cell division; that is, the approximation assumes that GFP is stable and the main mechanism for decreasing GFP per cell is dilution due to cell division rather than active degradation. So, instead of measuring *γ* directly, we used the cell growth rate 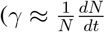, indicating how quickly the population is expanding per cell) to estimate it. This allows us to determine *γ*, which is then used in the expression 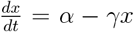 to calculate promoter activity *α*. Thus, from the measurements of cell density and fluorescence, the rate of change of cell density and fluorescence was calculated, and using these, the promoter activity was measured.

