## Supplementary for "Design and Characterisation of a Sensor Integrating Chemical and Temperature Inputs for Gene Expression"

Vinod Kumar and Shaunak Sen*

*Department of Electrical Engineering, Indian Institute of Technology Delhi, Hauz Khas, New Delhi - 110016, India.*

### Supplementary Figures

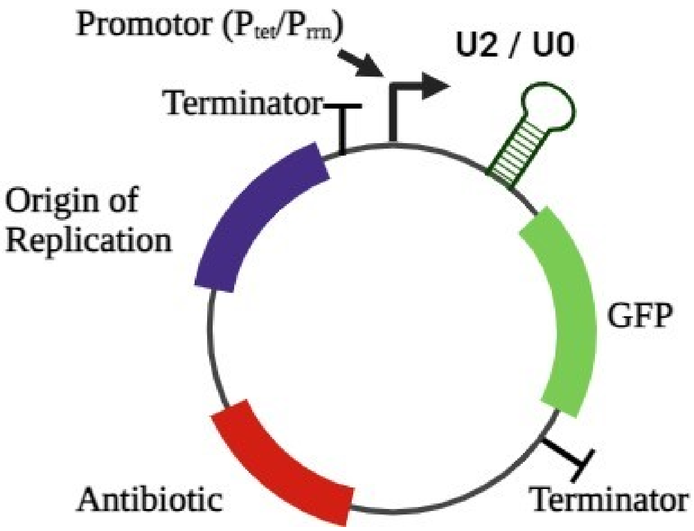


Figure 1: The plasmid schematic with necessary components used in this experimental study.

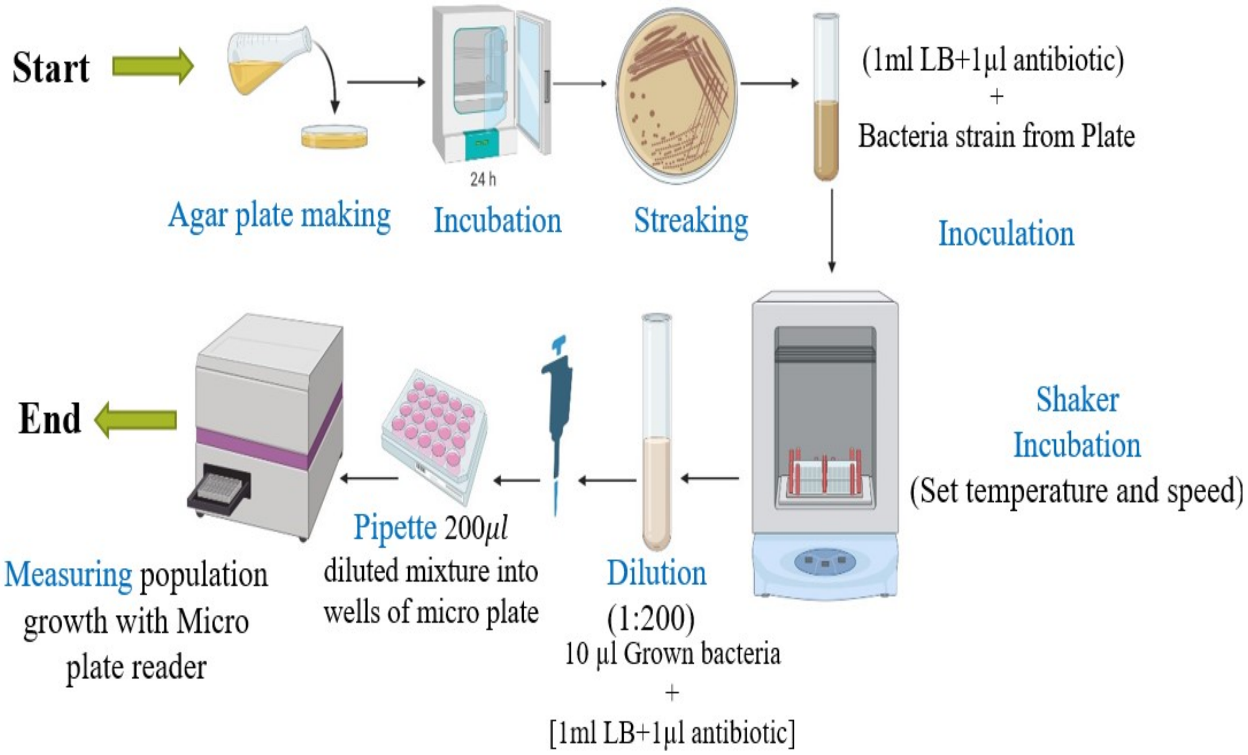


Figure 2: Material and procedure. Experiments are performed in vitro and start with transforming the plasmids into *E. coli* cells on the petri dishes (Agar plates), followed by an inoculation process in test tubes. The culture is grown optimally and sufficiently at 37 degrees at a set speed of 250 RPM in an incubator shaker for 12- 16 hours. This culture is then diluted and incubated for another 2-3 hours, which is then pipetted into wells of a transparent bottom microplate. This microplate is then inserted into a microplate reader, which measures fluorescence and optical density.

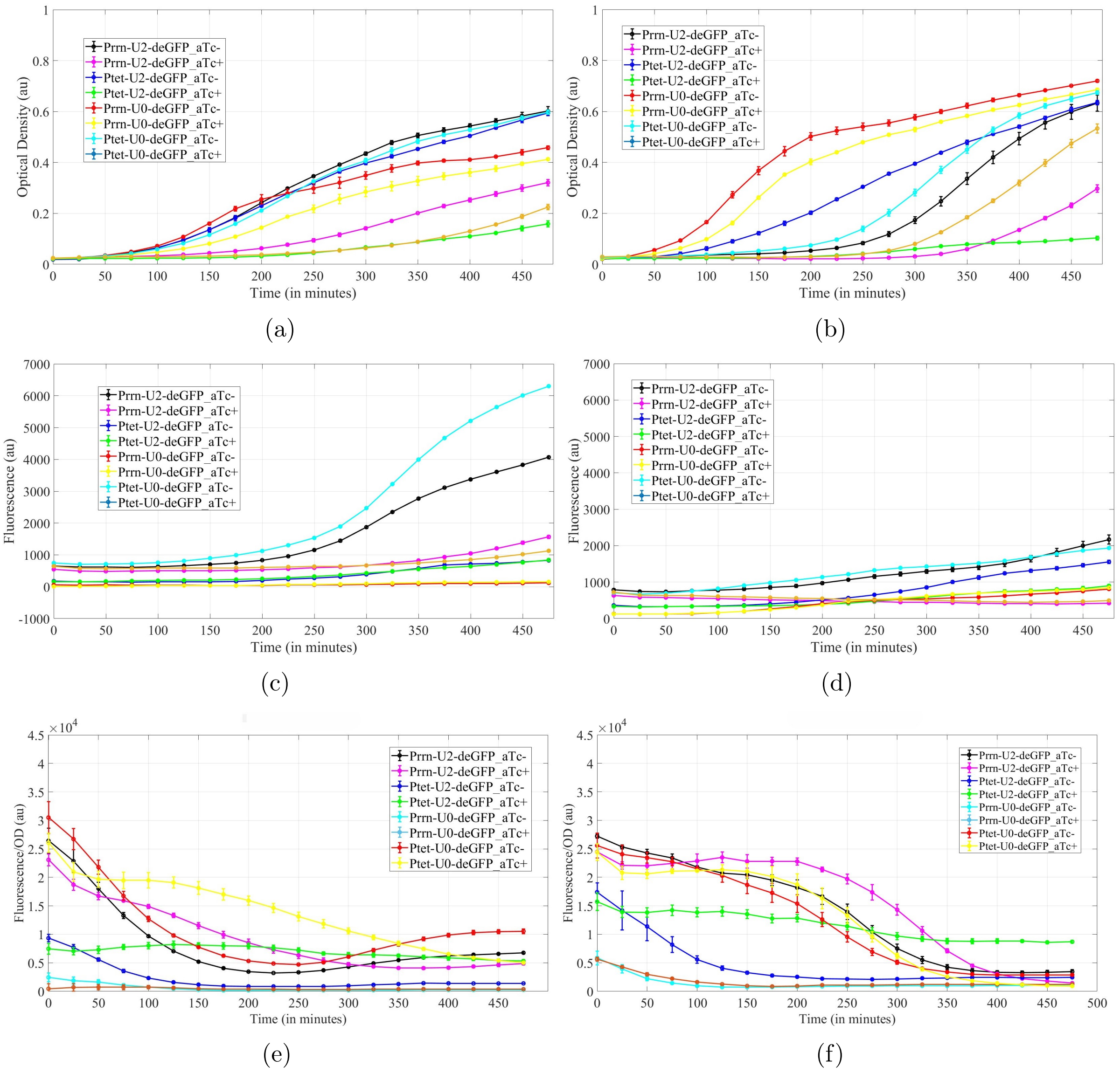


Figure 3: Optical Density (OD), Fluorescence (F), and F/OD for all constructs at (a),(c),(e) 29 °C, and (b),(d),(f) 37 °C.

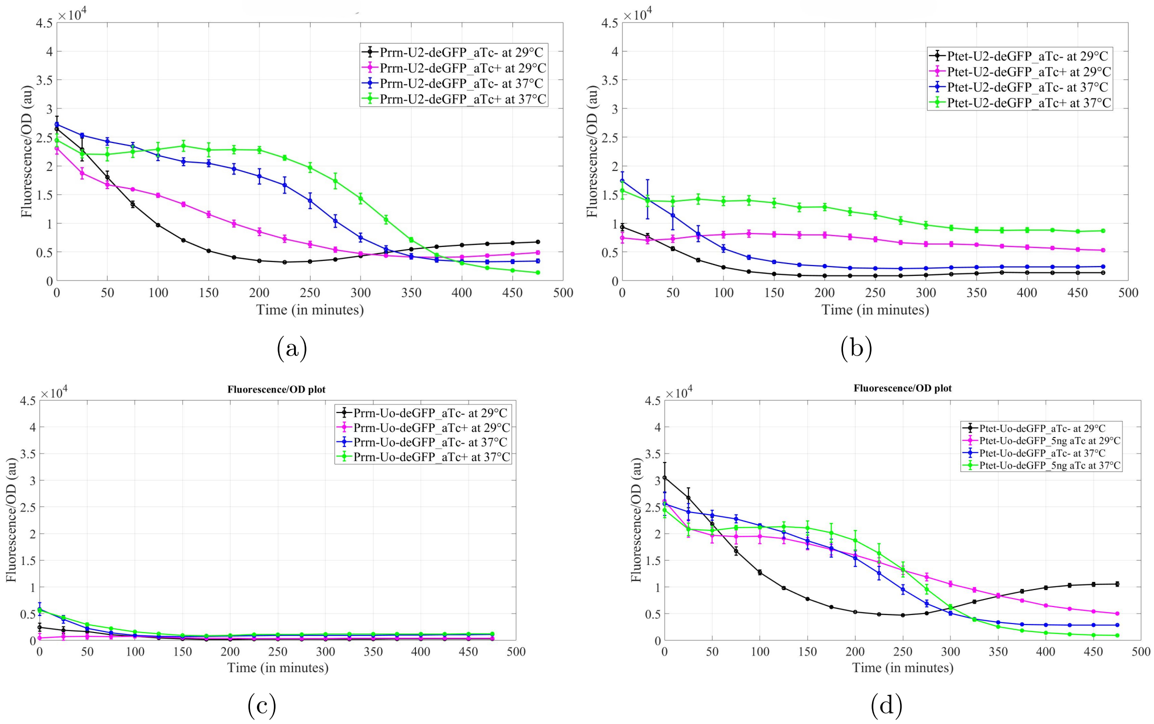


Figure 4: GFP/OD plots for constructs with, (a) Constitutive promoter $P_{rrn}$ and RNA thermometer($U2$), (b) Promoter $P_{tet}$ and RNA thermometer($U2$), (c) Consecutive and $U0$ (No thermometer or Weak thermometer, and (d) Promoter $P_{tet}$ and weak RNA thermometer ($U0$)) at 37 °C, and 29 °C.

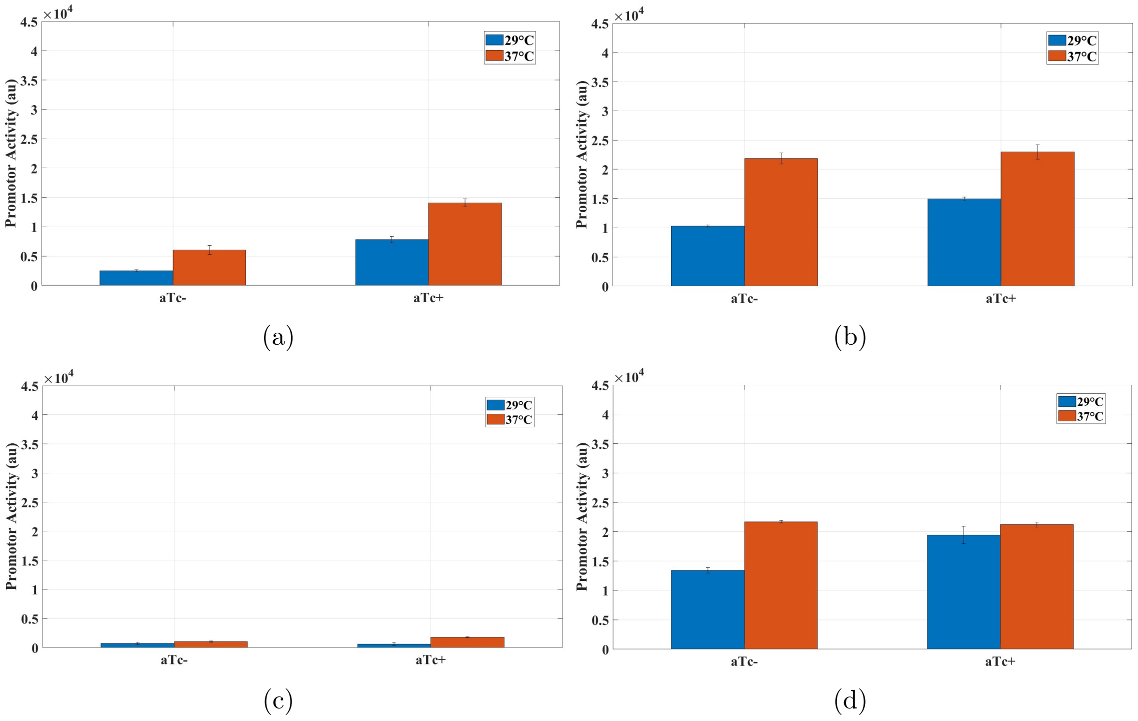


Figure 5: Promoter activity ($\alpha$) at the $100^{th}$ minute after the induction of aTc. Blue and Red bars indicate activities at 29 °C and 37 °C, respectively, for constructs with (a) Promoter $P_{tet}$ and RNA thermometer($U2$), (b) Constitutive promoter $P_{rrn}$ and RNA thermometer($U2$), (c) Consecutive and $U0$ (No thermometer or Weak thermometer, and (d) Promoter $P_{tet}$ and weak RNA thermometer ($U0$)).

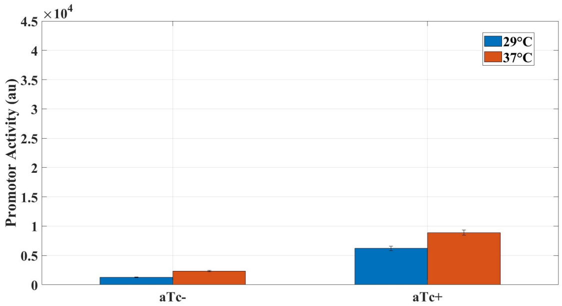


(a)

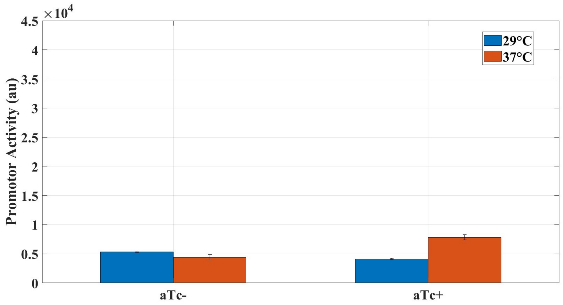


(b)

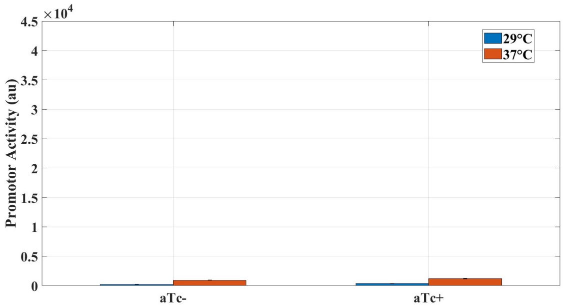


(c)

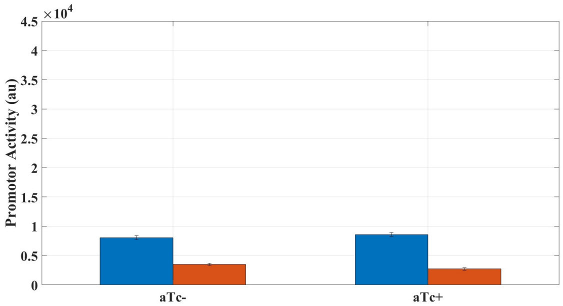


(d)

Figure 6: Promoter activity ($\alpha$) at the $350^{th}$ minute after the induction of aTc. Blue and Red bars indicate activities at 29 °C and 37 °C, respectively, for constructs with (a) Promoter $P_{tet}$ and RNA thermometer($U2$), (b) Constitutive promoter $P_{rrn}$ and RNA thermometer($U2$), (c) Consecutive and $U0$ (No thermometer or Weak thermometer, and (d) Promoter $P_{tet}$ and weak RNA thermometer ($U0$)).

### Parameter formulation: Asymmetry($a$) and Logic type($l$)

For characterizing the behavior of the combinatorial circuit, a 2-D logic symmetry space is used, which is defined by two parameters, logic type ($l$) and asymmetry ($a$), and plotted as ($l$, $a$) on this logic-symmetry space. '$l$' indicates if the behavior is towards more OR-like, more AND-like, or in between these two. '$a$' assesses the extent to which a construct's response favors one input (aTc or temperature) over another. A value of $a = 0$ signifies a symmetric response to both inputs (e.g., AND-like or OR-like behavior), whereas $a = 1$ denotes a single-input gate (SIG-like), where only one input dictates the response. The computation of the logic type ($l$) is based on the Per-read $X_{logic}$ strategy and the construct-wide logic behavior, whereas computation of asymmetry ($a$) is based on the absolute difference of effect of aTc and temperature, which shows how much one input effect dominates. The absolute difference is then normalized by the max $[E_{aTc}, E_{temp} + 0.01]$ to scale $a$ to $[0, 1]$. $\alpha_{\text{norm}}$ is the normalized $\alpha$ values, i.e, $\frac{\alpha}{15739}$ for each construct at the four input combinations: $[0, 0], [1, 0], [0, 1], [1, 1]$. The value 15739 is the basal value, which is the control's (constitutive) activity under standard growth conditions without aTc stimulation. The formulas used to compute both parameters are as follows.

**1. Asymmetry($a$):**

$$E_{\text{aTc}} = \alpha_{\text{norm}}c(2) + \alpha_{\text{norm}}c(4) \quad \text{(corresponding to inputs [1,0] and [1,1])}, \tag{1}$$

$$E_{\text{Temp}} = \alpha_{\text{norm}}c(3) + \alpha_{\text{norm}}c(4) \quad \text{(corresponding to inputs [0,1] and [1,1])}, \tag{2}$$

$$a = \frac{|E_{\text{aTc}} - E_{\text{Temp}}|}{\max(E_{\text{aTc}}, E_{\text{Temp}}) + 0.01} \tag{3}$$

Here, $E_{\text{aTc}}$ and $E_{\text{temp}}$ are the effects of aTc and temperature, respectively, and computed as,

$E_{\text{aTc}} =$ Sum of $\alpha_{\text{norm}}$ when aTc $= 1$ i.e when $[1, 0] + [1, 1]/c[2] + c[4]$.

$E_{\text{temp}} =$ Sum of $\alpha_{\text{norm}}$ when temp $= 1$ i.e when $[0, 1] + [1, 1]/c[3] + c[4]$.

The number 0.01 prevents division by zero if both effects are zero. This is interpreted as, if $E_{\text{aTc}} \approx E_{\text{temp}}$, than $a = 0$ (symmetric) more AND or OR like logic. If one input is much larger (e.g., $E_{\text{aTc}} >> E_{\text{temp}}$) than $a \approx 1$ (asymmetric or SIG like).

**Calculation:** $P_{tet} - U2 - deGFP$

$\alpha_{\text{norm}} = [0.053, 0.0139, 0.464, 0.747]$

$E_{\text{aTc}} = 0.139 + 0.747 = 0.866$

$E_{\text{temp}} = 0.464 + 0.747 = 1.211$

$a = \frac{|(0.866 - 1.211)|}{max(0.866, 1.211) + 0.01}$

$= \frac{0.325}{1.211 + 0.01} \approx 0.266$, this shows symmetry, which means both contribute almost the same.

**Conclusion:**$\Longrightarrow$ Low $a$ for AND$-$like ($P_{tet}-U2-deGFP$) and OR$-$like ($P_{tet}-U0-deGFP$) reflects balanced input effects, although $a$ is higher for $P_{tet} - U2 - deGFP$ indicating a more input-specific bias then $Ptet - U0 - deGFP$.

**2. Logic type ($l$):** $l = 0$ (More OR like), and $l = 1$ (More AND like)

$\alpha > 0.7$ (High activity), $\alpha < 0.465$ (Low activity) were chosen as thresholds. Considering these thresholds, $x_{logic}$ is considered 0 or 1 depending upon the activity of the combination of inputs,

**Calculation:** $P_{tet} - U2 - deGFP$

$\alpha_{\text{norm}} = [0.053, 0.0139, 0.464, 0.747]$

$[0, 0] : \alpha_{\text{norm}} = 0.053 < 0.465 \implies x_{logic} = 0$

$[1, 0] : \alpha_{\text{norm}} = 0.0139 < 0.465 \implies x_{logic} = 0$

$[0, 1] : \alpha_{\text{norm}} = 0.464 < 0.465 \implies x_{logic} = 0$

$[1, 1] : \alpha_{\text{norm}} = 0.747 > 0.7 \implies x_{logic} = 1$

**Conclusion:** $x_{logic} = [0, 0, 0, 1]$ (as per $x_{logic}$ construct 1 is seen to be AND-like logic gate in figure 2(b) in the manuscript)

Construct-wide $l$:

If $\alpha_{\text{norm}}[1, 1] > 0.7$ and $\alpha_{\text{norm}}[0, 0], [1, 0], [0, 1] < 0.465$ set $l = 1$

$\alpha_{\text{norm}}[1, 0], [0, 1], [1, 1] > 0.7$ and $\alpha_{\text{norm}}[0, 0] < 0.465$ set $l = 0$.

$\implies P_{tet} - X - deGFP : \alpha_{\text{norm}}[0.747] > 0.7$ other $[0.053, 0.139, 0.464] < 0.465$

$\implies l = 1 \implies$ High activity only when both inputs are ON (**AND-like**).

Similarly,

$\implies P_{tet} - U0 - deGFP : \alpha_{\text{norm}}[0.301, 0.727, 0.878, 0.962] > 0.7$, and $[0, 0] < 0.465$ (**OR-like**).

For other patterns (complex/additive), we computed the average logic score of intermediate reads $[0, 1], [1, 0]$. $l =$ mean $(x_{logic}[1, 0], x_{logic}[0, 1])$. For intermediate reads/inputs, we determined whether the construct's response ($\alpha_{\text{norm}}$) is closer to OR-like or AND-like. The interpolation formula thus provides a smooth transition between these extremes (AND or OR) when $\alpha_{\text{norm}}$ is neither clearly low nor clearly high.
